# Patterns of Z chromosome divergence among *Heliconius* species highlight the importance of historical demography

**DOI:** 10.1101/222430

**Authors:** Steven M. Van Belleghem, Margarita Baquero, Riccardo Papa, Camilo Salazar, W. Owen McMillan, Brian A. Counterman, Chris D. Jiggins, Simon H. Martin

## Abstract

Sex chromosomes are disproportionately involved in reproductive isolation and adaptation. In support of such a ‘large-X’ effect, genome scans between recently diverged populations or species pairs often identify distinct patterns of divergence on the sex chromosome compared to autosomes. When measures of divergence between populations are higher on the sex chromosome compared to autosomes, such patterns could be interpreted as evidence for faster divergence on the sex chromosome, i.e. ‘faster-X’, or barriers to gene flow on the sex chromosome. However, demographic changes can strongly skew divergence estimates and are not always taken into consideration. We used 224 whole genome sequences representing 36 populations from two *Heliconius* butterfly clades (*H. erato* and *H. melpomene*) to explore patterns of Z chromosome divergence. We show that increased divergence compared to equilibrium expectations can in many cases be explained by demographic change. Among *Heliconius erato* populations, for instance, population size increase in the ancestral population can explain increased absolute divergence measures on the Z chromosome compared to the autosomes, as a result of increased ancestral Z chromosome genetic diversity. Nonetheless, we do identify increased divergence on the Z chromosome relative to the autosomes in parapatric or sympatric species comparisons that imply post-zygotic reproductive barriers. Using simulations, we show that this is consistent with reduced gene flow on the Z chromosome, perhaps due to greater accumulation of species incompatibilities. Our work demonstrates the importance of constructing an appropriate demographic null model in order to interpret patterns of divergence on the Z chromosome, but nonetheless provides evidence to support the Z chromosome as a strong barrier to gene flow in incipient *Heliconius* butterfly species.

## Introduction

Comparisons between genomes of diverging populations or species have revealed elevated differentiation on the sex chromosomes in several animals, such as fly-catchers (Ellegren *et al*. 2012), crows (Poelstra *et al*. 2014), Darwin’s finches (Lamichhaney *et al*. 2015), ducks (Lavretsky *et al*. 2015) and *Heliconius* butterflies (Kronforst *et al*. 2013; Martin *et al*. 2013; Van Belleghem *et al*. 2017). These patterns of elevated sex chromosome divergence are sometimes readily interpreted as the result of increased reproductive isolation and reduced admixture on the sex chromosomes and, thus, ascribed to a large-X effect (BOX 1). However, it remains unresolved whether such elevated sex-linked divergence actually results from more rapid accumulation of isolating barriers on the sex chromosome, or could be explained by differences in effective population size between the sex chromosomes and the autosomes (Pool & Nielsen 2007; Meisel & Connallon 2013; Wolf & Ellegren 2017).

#### BOX 1. Consequences of hemizygous sex chromosomes

##### Large-X (or Z) effect and what can cause it

Sex chromosomes have been repeatedly shown to have a disproportionate role during speciation (Coyne & Orr 2004), demonstrated by three widespread intrinsic postmating effects (Turelli & Moyle 2007; Johnson & Lachance 2012); (i) Haldane’s rule, (ii) reciprocal-cross asymmetry of hybrid viability and sterility, and (iii) the large-X effect. *Haldane’s rule* states that where only one sex of the hybrids has reduced viability or fertility, that sex is most commonly the heterogametic sex (Haldane 1922). *Asymmetry of hybrid viability and sterility* refers to the situation where reciprocal crosses often differ in their incompatibility phenotype (Turelli & Moyle 2007). Finally, the *large X-effect* highlights the disproportionate contribution of the sex chromosomes to the heterogametic and asymmetric hybrid inviability/sterility in backcross families (Coyne & Orr 1989).

Haldane’s rule can generally be explained by between-locus ‘Bateson-Dobzhansky-Muller incompatibilities’ (BDMs) (Dobzhansky 1935; Muller 1942; Orr 1996), in which divergent alleles at different loci become fixed between populations and cause inappropriate epistatic interactions only when brought together in novel hybrid allele combinations (Coyne & Orr 2004). If interacting loci include recessive alleles on the sex chromosome, only the heterogametic sex will suffer incompatibilities (Turelli & Orr 1995). Additionally, BDMs between autosomal loci and the sex chromosomes can be specific to a particular direction of hybridization due to their uniparental inheritance and, thus, also explain asymmetric reproductive isolation (Turelli & Moyle 2007). Hence, hemizygous expression of recessive alleles on the sex chromosome has been put forward as a trivial cause for the disproportionate role of the sex chromosomes during speciation (dominance theory) (Turelli & Orr 1995).

In contrast to the dominance theory, there is however a large body of observations and theory that propose alternative or additional explanations that can cause a large-X effect (Wu & Davis 1993; Presgraves 2008). These factors include faster male evolution resulting from intense sexual selection among males (Wu & Davis 1993), meiotic drive (Frank 1991), dosage compensation (Jablonka & Lamb 1991) and faster-X evolution (Charlesworth *et al*. 1987; Vicoso & Charlesworth 2006). Faster-X evolution is predicted if adaptive new mutations are on average partially recessive (Charlesworth *et al*. 1987). In *Drosophila*, faster-X evolution has been studied extensively. Although it is not ubiquitous, there is clear evidence for faster-X divergence and adaptation (Counterman *et al*. 2004), particularly for X-linked genes expressed in male reproductive tissues (reviewed in Meisel & Connallon 2013). In Lepidoptera (butterflies and moths), Haldane’s rule and the large X (or Z) effect have been reported for numerous species, yet the females are the heterogametic sex (Sperling 1994; Prowell 1998; Presgraves 2002). Since lepidopteran females are heterogametic, faster male evolution is insufficient to explain Haldane’s rule and the large X effect, but faster-X evolution remains a viable explanation (Sackton *et al*. 2014). Moreover, in Lepidoptera, the large Z effect extends beyond intrinsic isolating barriers and there are differences in many traits and behaviors that map disproportionately to the Z chromosome (Sperling 1994; Prowell 1998). These observations are consistent with the faster accumulation of differences on the Z chromosome (faster-X evolution).

##### Factors affecting sex/autosome diversity ratios

Apart from population size changes, factors that can result in deviations from the expected three-quarter X/autosome (X/A) diversity ratio, and could thus potentially affect divergence measures, include (i) sex-biased demographic events leading to different effective population sizes of males and females (Charlesworth 2001), (ii) selective sweeps and background selection differently affecting the sex chromosomes (Charlesworth 2012) and (iii) differences in mutation rates between sexes or between the sex chromosomes and the autosomes (Sayres & Makova 2011; Johnson & Lachance 2012).

First, different population sizes of males and females can influence the X/A diversity ratio because two-third of the X chromosome population is transmitted through females. A male biased population would thus decrease the X/A diversity ratio below three-quarters, whereas a female biased population would increase the ratio. This effect would be opposite in female heterogametic sex systems (ZW).

Second, the hemizygous expression of the sex chromosome could result in both higher purifying selection and more efficient selection of beneficial recessive mutations (~selective sweeps) and result in a decrease of the expected X/A diversity ratio (Charlesworth *et al*. 1987). Additionally, differences in recombination rates can lead to different extent of loss of variation through linked selection and thus background selection (Charlesworth 2012). In Lepidoptera, meiosis is commonly achiasmatic (no recombination) in the heterogametic sex (females) (Suomalainen *et al*. 1973; Turner & Sheppard 1975). A reduction in recombination rate on the sex chromosomes compared to autosomes, that are commonly found in *Drosophila* (Vicoso & Charlesworth 2009), should thus not be expected to decrease Z/A diversity ratios through increased background selection in *Heliconius*. On the other hand, it has been suggested that effective recombination should be higher, and thus background selection lower, for the Z chromosome when recombination is absent in females (Charlesworth 2012). This is because the Z chromosomes spend two-third of their time in recombining males, whereas autosomes only spend half of their time in recombining males.

Third, because the male germ line generally involves more cell divisions and thus opportunities for replication errors, sex-linked genes may have different mutation rates. Because X-linked genes spend only one-third of their time in males and two-thirds of their time in females, the X chromosome may be subjected to a lower mutation rate. Conversely, the Z chromosome spends two-third of its time in males and may therefore become enriched in genetic variation compared to the autosomes (Vicoso & Charlesworth 2006; Sayres & Makova 2011; Johnson & Lachance 2012). Such increased mutation rates on the Z chromosome could also increase the rate of divergence between Z chromosomes (Kirkpatrick & Hall 2004).

When comparing divergence between genomic regions, such as sex chromosomes versus autosomes, measures of population divergence are influenced by within population diversity (Charlesworth 1998; Cruickshank & Hahn 2014) (BOX 2). This is explicitly the case for relative measures such as *F_ST_
*, but also influences absolute measures of divergence such as *d_XY_
*. For absolute divergence measures, this is because the genetic divergence between two alleles sampled from two species includes both divergence accumulated post-speciation, but also diversity already present in the ancestral population before the split. The latter is strongly dependent on effective population size. In a population under equilibrium conditions where the two sexes have an identical distribution of offspring number, the X chromosome effective population size and genetic diversity is expected to be three-quarters that of the autosomes. Deviations from this ratio can result from multiple unique features of the sex chromosomes (BOX 1), and population size changes in particular can have strong differential influence on sex chromosome compared to autosomal diversity (Pool & Nielsen 2007). Previous studies attempted to control for differences in effective population size on the sex chromosome, for instance among recently diverged duck species from Mexico, but such studies generally do not account for population size changes (Lavretsky *et al*. 2015). In order to interpret both relative and absolute measures of divergence on the sex chromosomes as evidence of a disproportionate contribution to species divergence and/or reduced admixture, we need to also account for demographic changes that can influence diversity of the sex chromosomes.

#### BOX 2. Measures of divergence depend on population size

The mutational diversity in present-day samples is directly related to population size, structure and age. This diversity within populations directly impacts the rate of coalesce between populations (Figure B 1). This relationship can be seen with *F_ST_
*, which was developed to measure the normalized difference in allele frequencies between populations (Wright 1931). The dependence of *F_ST_
* on population size can be understood by interpreting *F_ST_
* as the rate of coalescence within populations compared to the overall coalescence rate (Slatkin & Voelm 1991). Comparing pairs of populations with different effective population sizes will therefore show distinct *F_ST_
* estimates even when the split time is the same (Charlesworth 1998). Absolute divergence *d_XY_
* is the average number of pairwise differences between sequences sampled from two populations (Nei & Li 1979). *d_XY_
* is not influenced by changes to within population diversity that occur after the split, but does depend on diversity that was present at the time the populations split (Gillespie & Langley 1979). Therefore, population pairs that had a smaller population size at the time they split will, consequently, have smaller *d_XY_
* estimates. To compare pairs of populations that had different ancestral population sizes, *d_a_
* has been proposed, which subtracts an estimate of the diversity in the ancestral population from the absolute divergence measure *d_XY_
* (Nei & Li 1979). An approximation of ancestral diversity can be obtained by taking the average of the nucleotide diversity observed in the two present day populations. Such a correction should result in the ‘net’ nucleotide differences that have accumulated since the time of split.

**Figure B 1.**
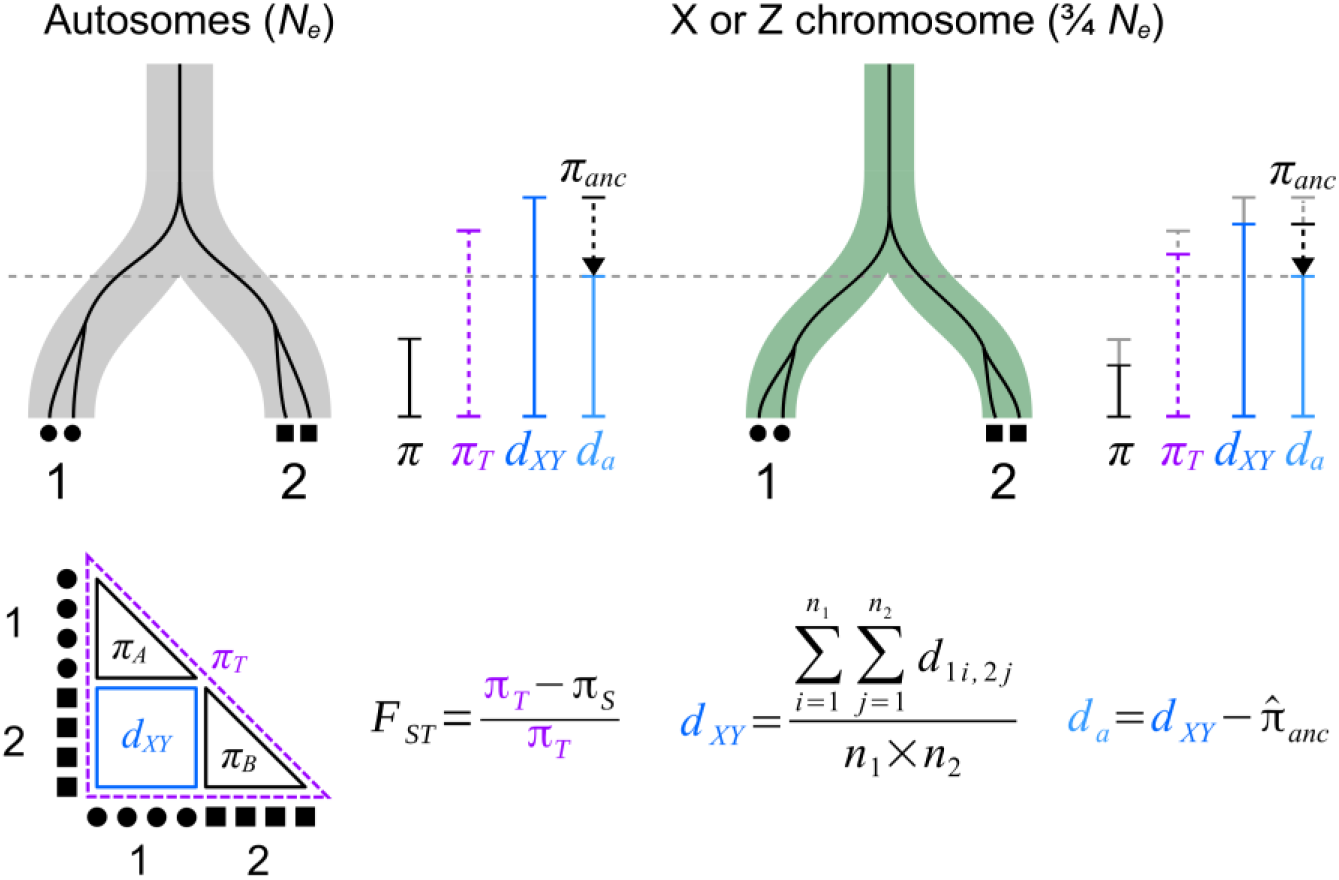
The effect of population size on the coalescent and measures of diversity and divergence. The branches represent two populations 1 and 2 that have split at a certain time (gray dashed line) and which can have a different size, such as the autosomes (gray) and X chromosome (green). The black lines show the coalescent of two alleles in each population. This coalescence process is influenced by the split time as well as the population size. Population size affects the nucleotide diversity within each population (*π*), the total nucleotide diversity (*π_T_
*) and absolute divergence *d_XY_
*, but not *d_a_
* as indicated by the vertical colored lines. For *d_a_, π_S_
* is used as the estimate of *π_anc_
*. The influence of population size on *F_ST_
* can be seen as resulting from a decrease in the denominator (*π_T_
*), but not in the numerator (*π_T_
* and *π_s_
* change proportionately).

To show how these different divergence measures perform, we simulated a simplified population split with varying degrees of migration (*m*) (Figure B 2). As expected, the values *F_ST_, d_XY_
* and *d_a_
* all increase when migration between populations decreases. *F_ST_
* and *d_XY_
* are clearly influenced by population size. While for *d_XY_
* this simply results from the variation present at the time of the split, *F_ST_
* does not show a simple linear relationship with population size. Only *d_a_
* represents the net accumulation of differences that can be compared between populations of different sizes, such as the sex chromosomes versus autosomes.

**Figure B 2.**
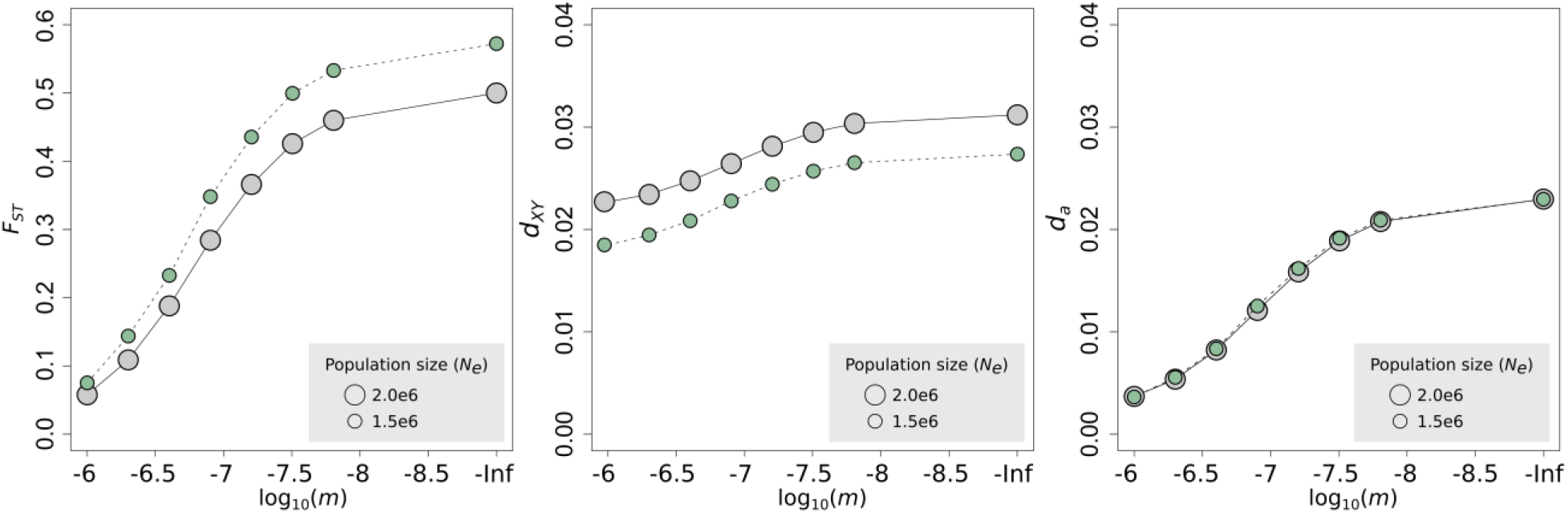
Simulated effect of population size differences on divergence measures *F_ST_, d_XY_
* and *d_a_
*. Simulations were performed for two populations that split 4 million generations ago and vary in their degree of migration (*m*). A lower effective population size, such as for the X chromosome (green) compared to autosomes (gray), results in higher *F_ST_
* and lower *d_XY_
* estimates, but has no effect on *d_a_
* under these assumptions.

Here, we explore diversity and divergence on the Z chromosome relative to the autosomes among populations of the *Heliconius erato* and *Heliconius melpomene* butterfly clades, using these different measures. The *H. erato* and *H. melpomene* clades represent unpalatable and warningly colored butterflies that have independently radiated into many divergent geographic races and reproductively isolated species. Within both clades, speciation has been accompanied by shifts in Müllerian mimicry (Mallet, McMillan, *et al*. 1998) and where populations come into contact, hybrid phenotypes usually have reduced survival rates due to strong frequency dependent selection against intermediate color pattern phenotypes (Mallet & Barton 1989; Jiggins *et al*. 1996; Naisbit *et al*. 2001; Merrill *et al*. 2012). Two species, *H. himera* and *H. e. chestertonii*, are geographic replacements of *H. erato* in dry Andean valleys. They are partially reproductively isolated, but individuals of hybrid ancestry make up about 10% of the population in narrow transition zones between forms (McMillan *et al*. 1997; Muñoz *et al*. 2010; Merrill *et al*. 2014). Similarly, *H. cydno* and *H. timareta* are geographic replacements of each other and both are broadly sympatric with *H. melpomene*. Here, both species are reproductively isolated from *H. melpomene* by a combination of pre- and post-mating isolation (Merot *et al*. 2017). Species integrity does not seem to involve structural variation such as chromosomal inversions (Davey *et al*. 2017). Instead, reproductive barriers include strong selection against hybrids, mate choice and post-zygotic incompatibilities (Figure 1A). Assortative mating has evolved in both the *H. erato* and *H. melpomene* clades (McMillan *et al*. 1997; Jiggins, Naisbit, *et al*. 2001; Muñoz *et al*. 2010; Merrill *et al*. 2014). In the *H. erato* clade, sterility and reciprocal-cross asymmetry of hybrid sterility has been reported in crosses between *H. erato* and *H. e. chestertonii* (Muñoz *et al*. 2010), but hybrid sterility is absent between *H. erato* and *H. himera* (McMillan et al. 1997). In the *H. melpomene* clade, female sterility (Haldane’s rule) and reciprocal-cross asymmetry of hybrid sterility occurs in crosses between *H. melpomene* and *H. cydno* (Naisbit *et al*. 2002), *H. melpomene* and *H. heurippa* (Salazar *et al*. 2005) and *H. melpomene* and *H. timareta* (Sanchez *et al*. 2015), as well as between allopatric *H. melpomene* populations from French Guiana and those from Panama and Colombia (Jiggins, Linares, *et al*. 2001). In support of a large-X effect, sterility in these crosses (*H. melpomene* x *H. cydno, H. melpomene* x *H. heurippa* and *H. melpomene* x *H. timareta*) was found to be Z-linked.

**Figure 1.**
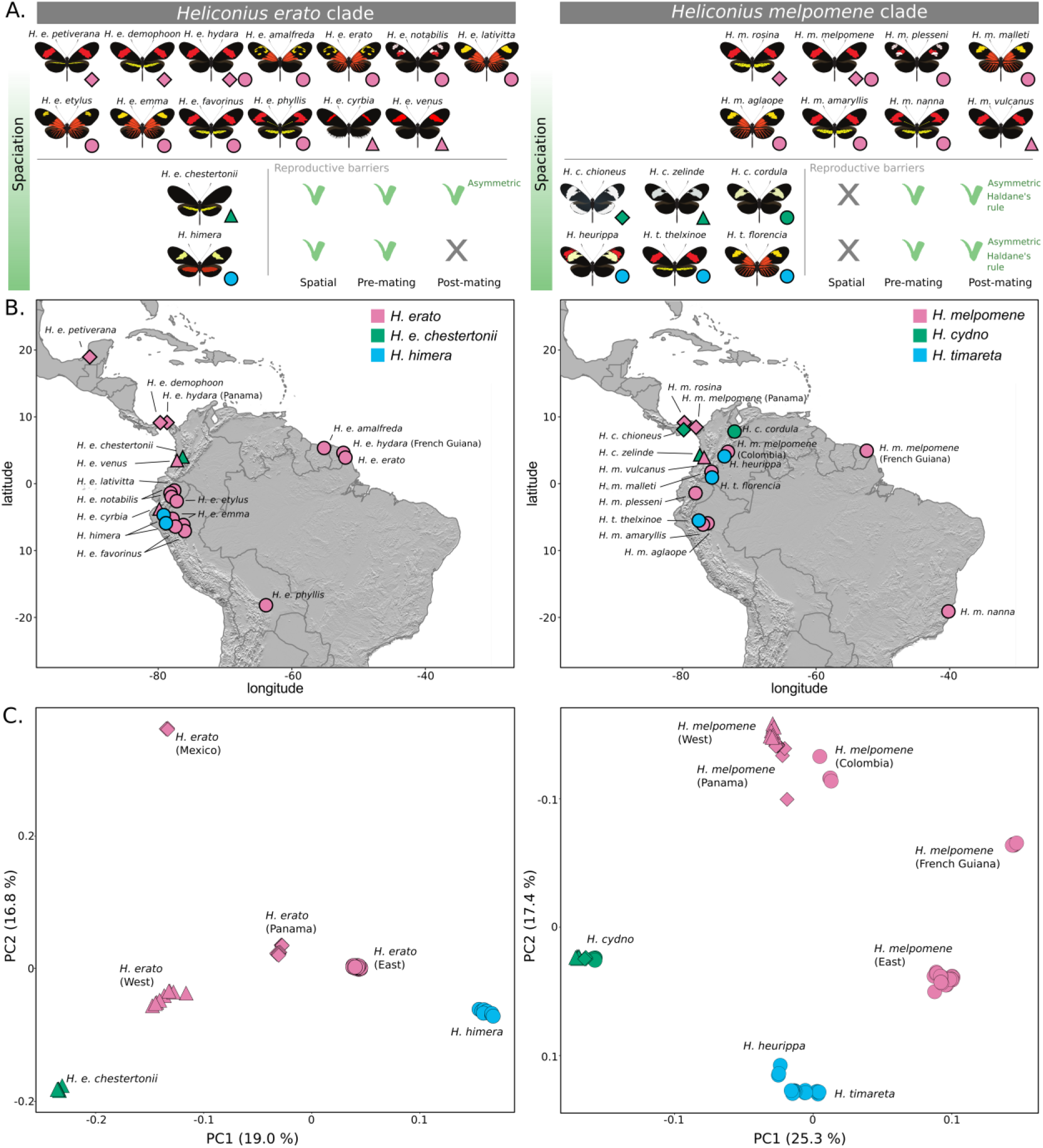
Speciation in the *Heliconius erato* and *Heliconius melpomene* clade and sampling. **(A.)**
*Heliconius erato chestertonii* (green) is reproductively isolated from *H. erato* (pink) by spatial separation (parapatry), mate choice and (asymmetric) reduced hybrid fertility of both sexes (i.e. no Haldane’s rule). *Heliconius himera* (blue) is reproductively isolated from *H. erato* by spatial separation and mate choice, but hybrids show no reduced fertility. *Heliconius cydno* (green) and *H. timareta* (blue) occur sympatrically with *H. melpomene* (pink) populations, but are both reproductively isolated by strong mate choice and (asymmetric) reduced fertility of heterozygous hybrids (i.e. Haldane’s rule). **(B.)** Localities of sampled populations included in this study. Within *H. erato, H. melpomene, H. timareta* and *H. cydno* names represent different races that display distinct color patterns. Shapes represent geographic regions; Mexico and Panama (diamond), west of the Andes (triangles) and east of the Andes (circles). **(C.)** PCA plots of autosomal SNP variation. Note that *H. m. nanna* has not been included in the PCA analysis as the signal of geographic isolation between *H. m. nanna* and the other populations dominates the signal (see Figure S1).

The presence of incipient species pairs with different levels of reproductive isolation allows us to examine the relative rate of autosomal and Z chromosomal evolution and the factors that are likely influencing patterns of divergence. We take advantage of a large genomic dataset composed of 224 whole genomes representing 20 populations of the *H. erato* clade and 16 populations of the *H. melpomene* clade. We also use simulations to evaluate the effect that demographic changes have on the estimate of relative rates of divergence on the Z versus the autosomes and demonstrate that in many comparisons demography can explain much of the observed elevated divergence on the Z relative to the autosomes. However, by taking into account geographic distance or autosomal divergence as a proxy for gene flow, we show that there is evidence for increased divergence on the Z chromosome for species pairs with known post-zygotic reproductive barriers. These rates of increased divergence likely reflect reduced admixture on the Z chromosome and provide support for the Z chromosome being a greater barrier to gene flow in some incipient *Heliconius* butterfly species.

## Results and discussion

### Population structure in Heliconius erato and Heliconius melpomene

We mapped a total of 109 *Heliconius erato* clade resequenced genomes to the *Heliconius erato* v1 reference genome (Van Belleghem *et al*. 2017) and 115 *Heliconius melpomene* clade genomes to the *Heliconius melpomene* v2 reference genome (Davey *et al*. 2016). These samples represent 20 *H. erato* clade and 16 *H. melpomene* clade populations covering nearly the entire geographic distribution of these species groups (Figure 1A and B).

Principal components analysis (PCA) of the autosomal SNP variation, performed using Eigenstrat SmartPCA (Price *et al*. 2006), grouped the *H. erato* clade samples mainly according to geography, apart from *H. himera* individuals from Ecuador and northern Peru and *H. e. chestertonii* from Colombia (Figure 1C). Four main geographic groups were apparent: Mexico, Panama, populations west of the Andes and populations east of the Andes. *Heliconius erato* populations east of the Andes as far as 3000 km apart were tightly clustered in the PCA analysis. The separate grouping of *H. himera* and *H. e. chestertonii* individuals supports these populations as representing incipient species that maintain their integrity despite ample opportunity for hybridization and gene flow (Jiggins *et al*. 1996; McMillan *et al*. 1997; Arias *et al*. 2008). In the PCA, *H. himera* was more closely related to the *H. erato* populations east of the Andes, whereas *H. e. chestertonii* was more closely related to the West Andean populations.

PCA analysis of the *H. melpomene* clade grouped individuals from west of the Andes and Panama closely together, with *H. melpomene* from Colombia being most similar to this population pair (Figure 1C). *Heliconius melpomene* populations from east of the Andes further clustered in three distinct groups, largely in agreement with geographic distance; populations from the eastern slopes of the Andes, the French Guiana population and *H. m. nanna* from Brazil (Figure 1C and Figure S1). While phylogenetic reconstructions have suggested that *H. melpomene* and the *H. cydno/timareta* clades are reciprocally monophyletic (Dasmahapatra *et al*. 2012; Nadeau *et al*. 2013; Martin *et al*. 2013), such patterns are hard to interpret from the PCA and patterns of relatedness may be influenced by more recent admixture. Nevertheless, *H. cydno* and *H. timareta* clustered distinctly. *Heliconius cydno* formed a distinct cluster with little difference between samples from Panama, west or east of the Andes. *Heliconius timareta* grouped most closely with *H. heurippa*, consistent with previous analysis (Nadeau *et al*. 2013; Arias *et al*. 2014).

### Z chromosome divergence in Heliconius erato and Heliconius melpomene

To compare rates of divergence between the Z chromosome and autosomes, we calculated three measures that are commonly used to compare divergence between populations, *F_ST_, d_XY_
* and *d_a_
*, between incipient species and population pairs of *H. erato* and *H. melpomene* (Figure 2 and Figure S2-4). All three measures of sequence divergence are calculated from mutational diversity in the data, but they are each dependent on population size in different ways (BOX 2). In *Heliconius, F_ST_
* has been frequently used to identify regions in the genome under strong divergent selection (Nadeau *et al*. 2012; Martin *et al*. 2013; Van Belleghem *et al*. 2017). In comparisons between parapatric color pattern races of both *H. erato* and *H. melpomene*, sharp *F_ST_
* peaks are present near the major color pattern loci, suggesting both strong divergent selection and reduced gene flow (Figure S2). Additionally, increased *F_ST_
* values can be observed on the Z chromosome in comparisons between populations with assortative mating and hybrid inviability and sterility (Figure 2 and Figure S2). However, *F_ST_
* is influenced by effective population sizes (BOX 2). It is therefore problematic to obtain insights about selection or migration when comparing genomic regions with different effective population sizes, such as the Z chromosome and autosomes. Given equal numbers of breeding males and females, the Z chromosome is expected to have an effective population size three-quarters that of the autosomes. Smaller population size and the resulting lower nucleotide diversity on the Z chromosome may, therefore, partly explain inflated *F_ST_
* estimates on the Z chromosome.

In contrast, the *d_XY_
* values on the Z chromosome tend to be similar to or slightly lower than the average values on the autosomes in most species comparisons of the *H. erato* and *H. melpomene* clade (Figure 2 and Figure S3). Under equilibrium conditions, *d_XY_
* on the Z chromosome is expected to be three-quarters that of the autosomes at the time of the split. As time progresses and differences between populations accumulate, the proportion of the coalescent that is effected by the ancestral population size will become smaller and the ratio of *d_XY_
* on the Z to *d_XY_
* on the autosomes is expected to move towards one. However, estimating the exact split time is difficult and finding the expected absolute divergence for the Z chromosome compared to the autosomes is complicated (Patterson *et al*. 2006). Moreover, in contrast to the expectation that the ratio of *d_XY_
* will move towards one, *d_XY_
* on the Z chromosome is higher than on the autosomes for *H. e. cyrbia - H. himera* and *H. e. venus - H. e. chestertonii* comparisons (Figure 2).

Finally, by subtracting an estimate of diversity in the ancestral population from the absolute divergence measure d_XY_, known as *d_a_
* (Nei & Li 1979), we obtain an estimate of nucleotide differences that have accumulated since the time of split (BOX 2). The *d_a_
* estimates show significantly higher divergence on the Z chromosome in the comparisons *H. himera - H. e. cyrbia, H. e. venus - H. e. chestertonii*, and in *H. melpomene - H. cydno* and *H. melpomene - H. timareta* (Figure 2 and Figure S4). Overall, the increased *d_XY_
* in the *H. e. cyrbia* and *H. himera* and the *H. e. venus - H. e. chestertonii* comparisons and the higher *d_a_
* values on the Z chromosome relative to the autosomes appear to support a faster rate of divergence between *Heliconius* species pairs on Z chromosome.

**Figure 2.**
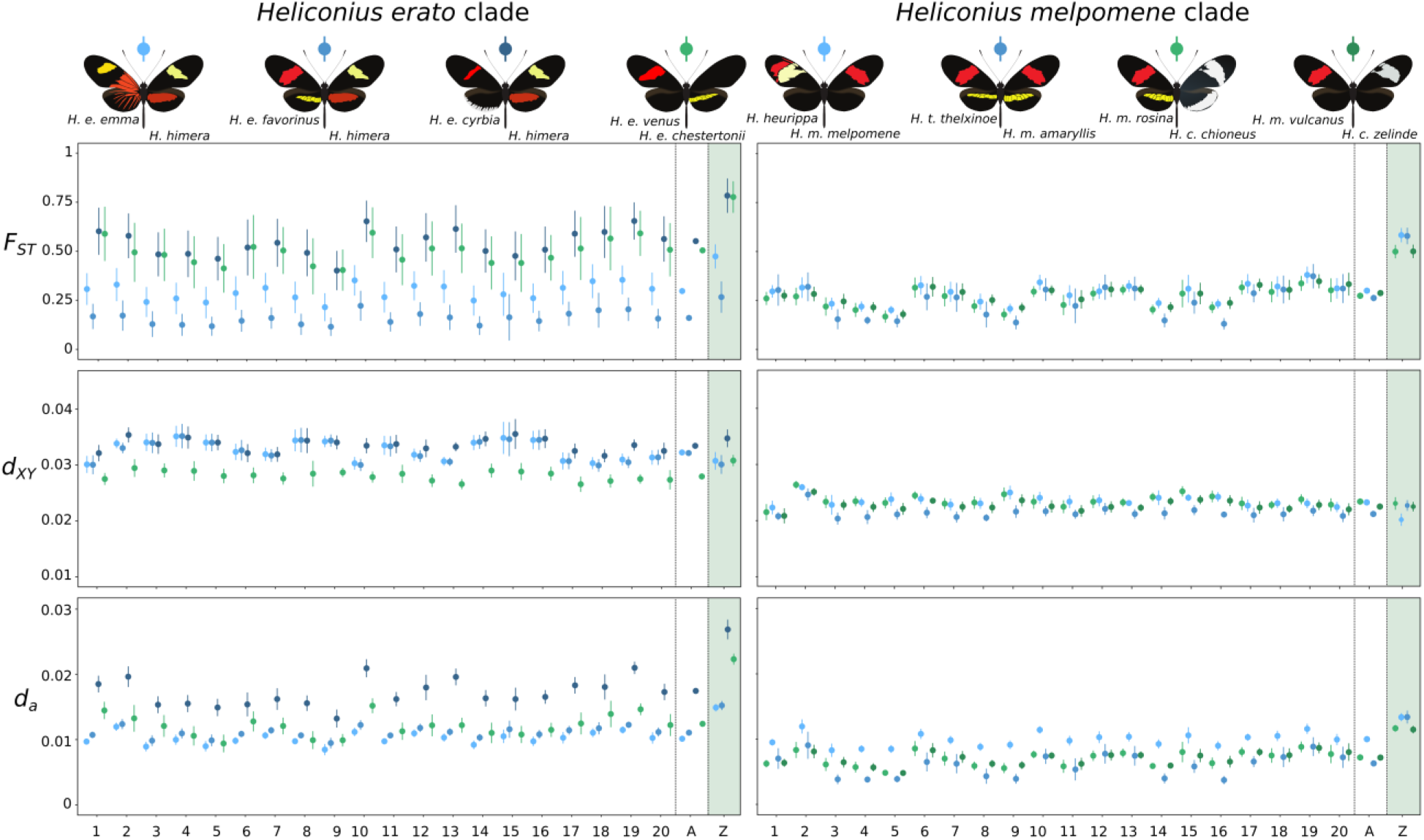
Averages of divergence measures at the autosomes (1-20) and Z chromosome in four parapatric/sympatric species comparisons of the *H. erato* and *H. melpomene* clade. Averages for each chromosome and 95 % confidence intervals (vertical bars) were estimated using block jackknifing using 1 Mb intervals. The vertical dashed lines highlight the averages over all autosomes (A) and the estimates for the Z chromosome. Values were calculated in 50 kb non-overlapping windows. See Figure S2, S3 and S4 for stepping window plots of *F_ST_, d_XY_
* and *d_a_
* values, respectively.

### Population size changes affect the Z chromosome differently

Apart from the overall difference in effective population size between the Z chromosome and autosomes, there are additional demographic factors that can contribute to differences in *F_ST_, d_XY_* and *d_a_* values between the Z chromosome and autosomes. Population size changes can alter the equilibrium expectation that Z-linked diversity should be three-quarters of autosomal diversity (Pool & Nielsen 2007). To explore this, we performed coalescent simulations of sequences from populations that underwent a single size change in the past, varying the time and magnitude of this event (Figure 3). In these simulated populations, the decrease in nucleotide diversity that follows population contraction occurs much faster than the increase in diversity that follows an expansion of the same magnitude (Figure 3A). This is because an increase in diversity requires mutation accumulation, whereas drift can rapidly remove variation to reach a new equilibrium. Additionally, population size changes have proportionately stronger effects on diversity on the Z chromosome compared to the autosomes (Figure 3B). This results from populations with a smaller effective population size, such as the Z chromosome, converging faster to their new equilibrium after a population size change (Pool & Nielsen 2007).

**Figure 3.**
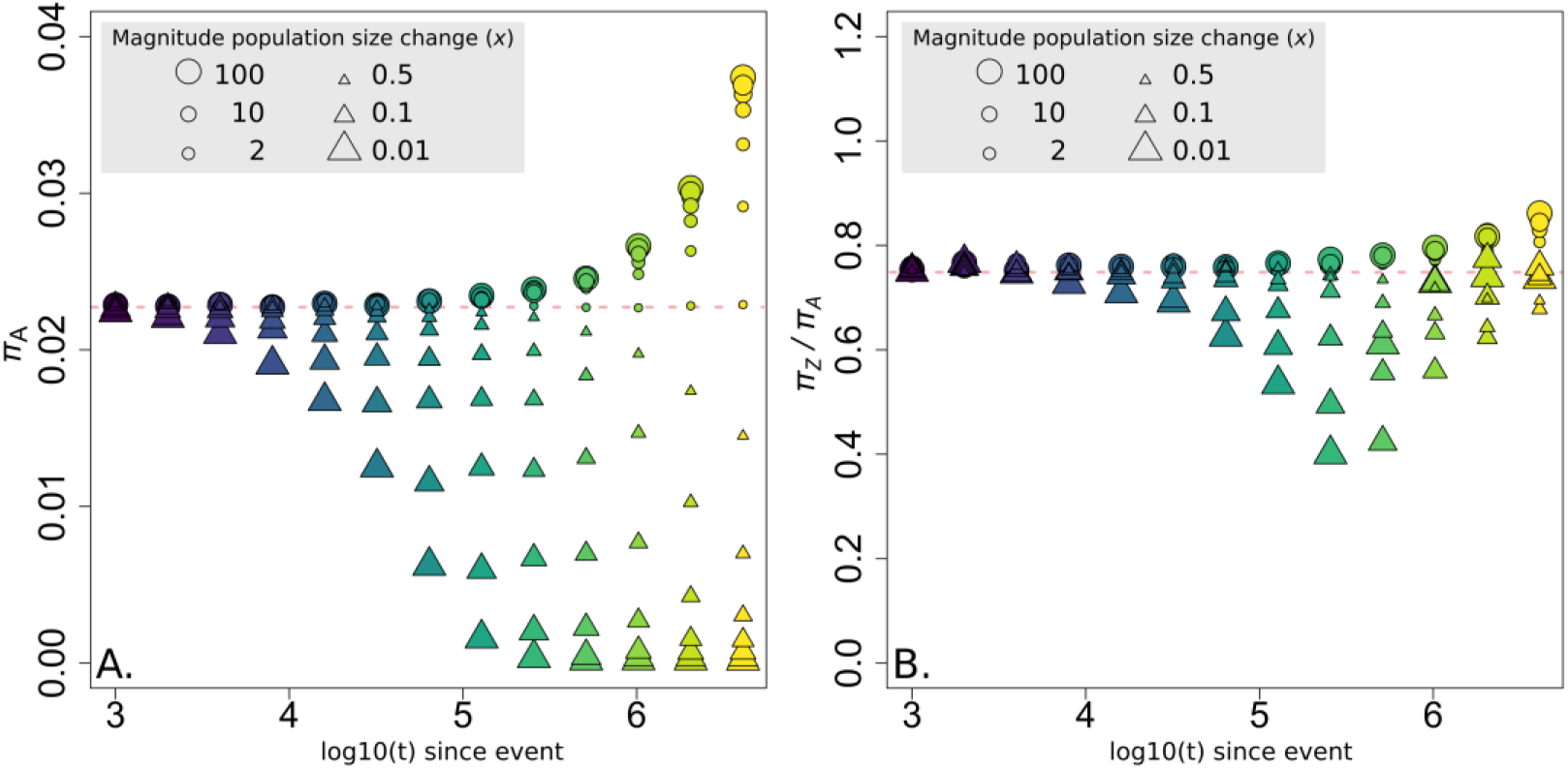
Simulated effect of population size change on nucleotide diversity on the autosomes (*π_A_
*) (A.) and the ratio of nucleotide diversity between the Z chromosome and autosomes (*π_Z_
*/*π_A_
*) (B.). Triangles indicate population size contractions, circles indicate population size increase. Colors and x-axis represent the time at which the population size change occurred in generations (log10(*t*)) with *N_e_
* equal to 3e6 and 2.25e6 for the autosomes and Z chromosome, respectively. The colors and time at which population size change occurred correspond to colors in the simulations in Figure 6. The pink dashed lines indicate expectations under neutrality.

The result is that divergence measures are differentially affected by population size change on the Z chromosome compared to the autosomes. The Z chromosome to autosome (Z/A) diversity ratio will be larger than expected in populations that experienced a recent expansion, and smaller than expected in those that experienced a recent contraction (Figure 3B). Therefore, in pairwise comparisons, if population size change occurred in the ancestral population before the two populations split, it would alter the ancestral Z/A diversity ratio and therefore confound comparisons of divergence between Z and autosomes using either relative or absolute measures of divergence, as all are influenced by ancestral diversity (Figure 4). By contrast if population size change occurred in one or both daughter populations after the split, it would affect the relative measures of divergence *F_ST_
* and *d_a_
*, but not absolute divergence (*dXY*), which is only dependent on ancestral diversity and not on diversity within each population. All the simulations were run for time scales relevant to *Heliconius* divergence and, therefore, demonstrate that a return to equilibrium values is unlikely after a population increase during the history of these species. Especially the effect of population size increases on Z/A diversity ratios can be long lasting during the evolutionary history of a population.

**Figure 4.**
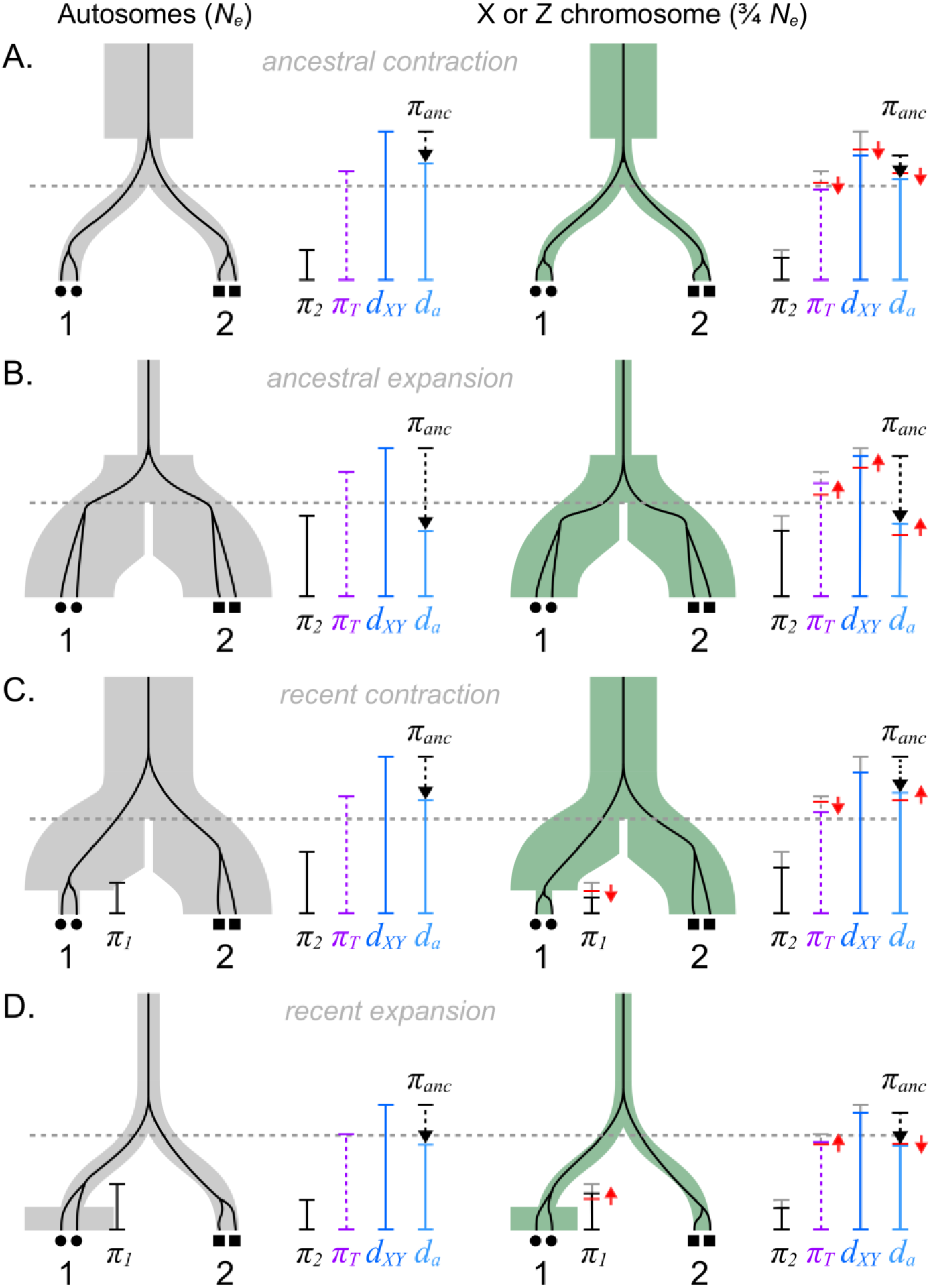
The effect of population size change on the coalescent and measures of diversity and divergence. The branches represent two populations 1 and 2 that have split at a certain time (gray dashed line) and which can have a different size, such as the autosomes (gray) and X chromosome (green). The black lines show the coalescent of two alleles in each population. This coalescence process is influenced by the split time as well as the population size (see BOX2). Moreover, red arrows show the disproportionate effect of population size change on diversity and divergence measures on the sex chromosome (or populations with a smaller size). Note that the effect size of a population size change will depend on the magnitude as well as the timing (see Figure 3). Exact expectations, including more complex size changes, can be calculated as demonstrated in Pool & Nielsen (2007).

### Demography and its influence on Z/A diversity ratios in Heliconius

To explore how population size changes might have affected Z/A diversity ratios and thus Z/A divergence comparisons within *Heliconius* clades, we used the behavior of Tajima’s *D* as a way to access likely population size changes within species. Tajima’s *D* is a population genetic measure commonly used to detect whether a DNA sequence is evolving neutrally (Tajima 1989). At a genome-wide scale, negative values reflect population size expansion, whereas positive values reflect population size decrease. Due to the different response of Tajima’s *D* to population size increase and decrease, Tajima’s *D* can give an indication of population size changes and their effect on nucleotide diversity. As the simulations show, negative Tajima’s *D* values (population size increase) are expected to correlate with increased nucleotide diversity, whereas positive Tajima’s *D* values (population size decrease) are expected to correlate with reduced nucleotide diversity (Figure 5). As with the Z/A diversity ratio, the timescale of the influence of population size change on Tajima’s *D* values is different for population expansion versus population contraction, as a population contraction returns to equilibrium much faster than an expansion. However, because smaller populations respond faster to such population size changes, the Tajima’s *D* values are also expected to be correlated with Z/A diversity ratios.

**Figure 5.**
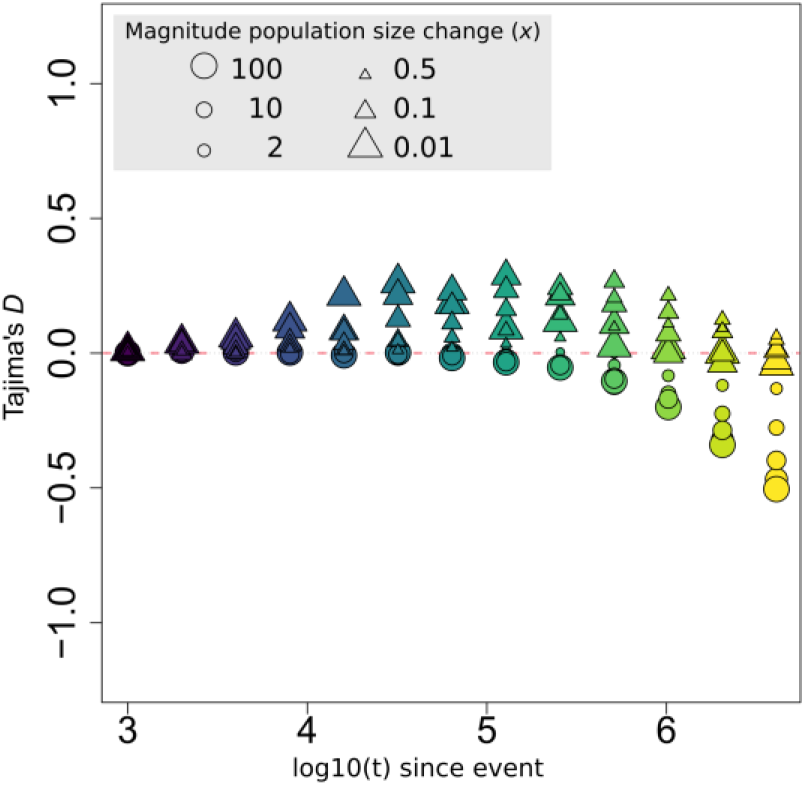
Simulated effect of population size change on Tajima’s *D* values. Triangles indicate population size contractions, circles indicate population size increase. Colors and x-axis represent the time at which the population size change occurred in generations (log10(*t*)) with *N_e_
* equal to 3e6 and 2.25e6 for the autosomes and Z chromosome, respectively. The colors and time at which population size change occurred correspond to colors in the simulations in Figure 6. The pink dashed line indicates expectations under neutrality.

Although this results in a complex relationship (Figure 6), *H. erato* clade populations that showed more negative Tajima’s *D* values (~population size increase) all had higher nucleotide diversity (*π*) values, as well as higher Z/A diversity ratios (Figure 6). There was a similar trend in the *H. melpomene* clade. It should be noted that multiple population size change events (e.g. population size expansion followed by a bottleneck) would further complicate the relation between Tajima’s *D* estimates and the expected nucleotide diversity as well as the Z/A diversity ratio. Nevertheless, the patterns among these *Heliconius* populations suggest that differences in diversity as well as differences in the Z/A diversity ratios are likely driven at least in part by population size changes. Given that samples assigned to a population were collected in close proximity, it is unlikely that estimated Tajima’s *D* values are influenced by hidden population structure in the data (which could result in negative Tajima’s *D* values).

**Figure 6.**
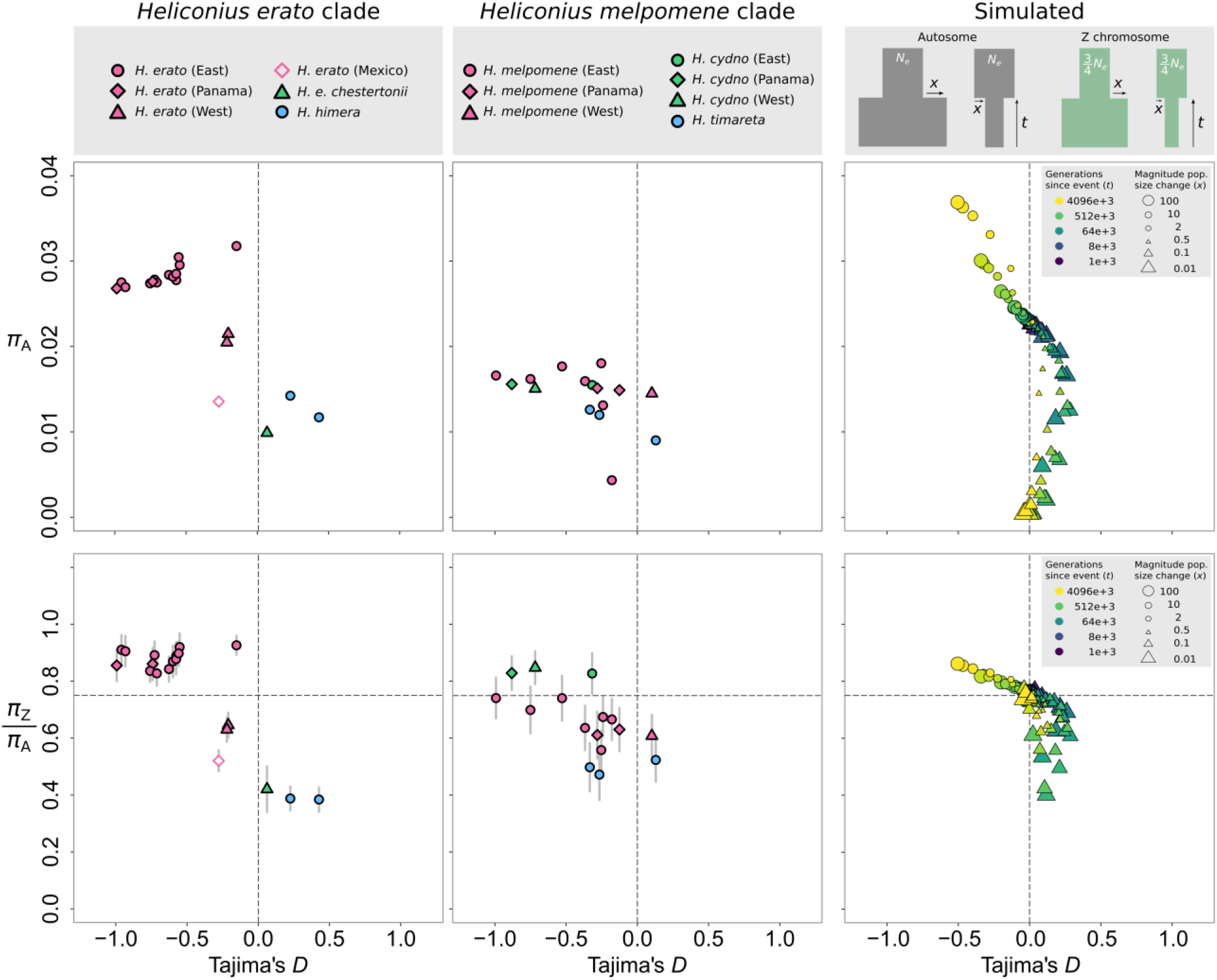
Relation between Tajima’s D, average nucleotide diversity on the autosomes (*π_A_
*) (upper panels) and the ratio of nucleotide diversity between the Z chromosome and autosomes (*π_Z_
*/*π_A_
*) (lower panels) for populations of the *H. erato* and *H. melpomene* clade and simulated data. Gray vertical bars represent 95% confidence intervals estimated from block-jackknifing, note that these are too small to see in the Tajima’s *D* versus *π_A_
* plots. Gray squares in the upper right panel represent the simulated population size changes. For the simulated data (right panels), the same data was used as in Figure 4 and Figure 5. Triangles indicate population size contractions, circles indicate population size increase, colors represent the time at which the population size change occurred in generations (*t*) with *N_e_
* equal to 3e6 and 2.25e6 for the autosomes and Z chromosome, respectively. The dashed lines indicate expectations under neutrality.

Broad patterns of nucleotide diversity can give some insight into the likely population size changes that have influenced our study species. Higher nucleotide diversity in *H. erato* clade as compared to the *H. melpomene* clade is consistent with the generally greater abundance of *H. erato* observed in nature (Mallet, Jiggins, *et al*. 1998) (Figure 6). These population size differences likely also explain the large differences in absolute divergence levels among the *H. erato* clade populations compared to the *H. melpomene* clade populations (Figure 7). Absolute divergence between *H. melpomene* and *H. timareta* or *H. cydno* is smaller than within population diversity of most *H. erato* populations (Figure 6), despite the former pairs being clearly distinct species.

**Figure 7.**
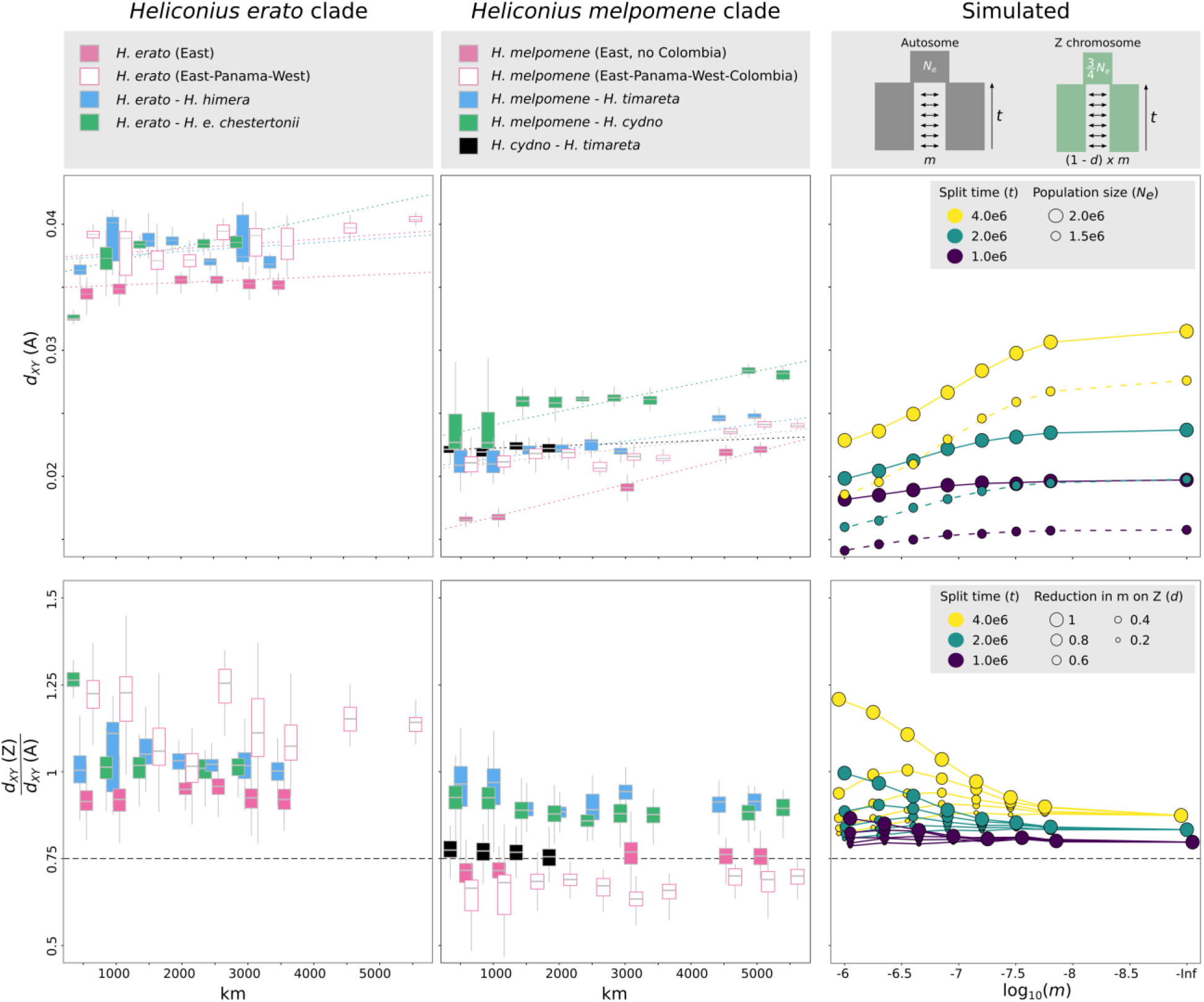
Relation between pairwise distance (km), autosomal *d_XY_
* (A) (upper panels) and the ratio of *d_XY_
* between the Z chromosome and autosomes (*d_XY_
*(Z)/*d_XY_
* (A)) for populations of the *H. erato* and *H. melpomene* clade and simulated data. Boxplots represent pairwise measures over 500 km bins. Gray squares in the upper right panel represent the simulated populations with different rates of migration (*m*) expressed as a proportion of the effective population size. For the simulated data (right panels), colors indicate split times of the two populations in generations (*t*) with *N_e_
* equal to 2e6 and 1.5e6 for the autosomes and Z chromosome, respectively. Size of circles indicates proportional reduction in migration on the Z chromosome.

While changes in population size can have strong effects on measures of sequence divergence, jointly considering patterns of variation on the Z chromosome and autosome can give further insights into the evolutionary history. Among *H. erato* populations from east of the Andes that show little differentiation, *d_XY_
*(Z)/*d_XY_
*(A) ratios are above 0.75 (0.91±0.11) (Figure 7) and there is also increased Z/A nucleotide diversity (Figure 6). This likely resulted from population size increase in the ancestral population. If this population size increase occurred before the divergence of *H. himera* and *H. e. chestertonii* from *H. erato* this could have contributed to elevated *d_XY_
*(Z)/*d_XY_
*(A) in these comparisons (Figure 6). The *d_XY_
*(Z)/*d_XY_
*(A) ratios among *H. melpomene* from east of the Andes are closer to the 0.75 ratio that would be expected immediately after the populations split (Figure 7). In contrast, while broader comparisons among *H. melpomene* populations east and west of the Andes, Panama and Colombia show clearly greater divergence, their *d_XY_
*(Z)/*d_XY_
*(A) ratios are much lower, consistent with a population size decrease deeper in the ancestry of *H. melpomene*. Finally, the lower *d_XY_
*(Z)/*d_XY_
*(A) ratios in *H. cydno - H. timareta* comparisons relative to the *H. melpomene* - *H. cydno* and *H. melpomene – H. timareta* comparisons suggests a population contraction of the ancestral population of *H. cydno* and *H. timareta*, but after they split from *H. melpomene*.

Within the *H. erato* clade, nucleotide diversity as well as Z/A diversity ratios were distinctly higher in populations from east of the Andes and Panama and lower in the *H. e. chestertonii* and *H. himera* populations (Figure 6). These populations shared a common ancestor, so differences in nucleotide diversity likely result from population size changes that occurred after divergence and thus confound the relative *F_ST_
* and *d_a_
* divergence measures. Although absolute divergence *d_XY_
* is clearly higher between *H. erato* and *H. himera* or *H. e. chestertonii* than among *H. erato* populations east of the Andes (Figure 7), a population size decrease in *H. himera* and *H. e. chestertonii* may inflate the *F_ST_
* and *d_a_
* estimates when comparing these populations to geographically abutting *H. erato* populations (Figure 2). Additionally, any population size changes that occurred before the split of *H. himera* from *H. erato* and *H. e. chestertonii* from *H. erato* may have affected current *dxY* estimates. Importantly, if such demographic changes differently affected the ancestor of *H. himera* as compared to the ancestor of *H. e. chestertonii*, the *d_XY_
* values may not necessarily reflect different degrees or stages of the speciation process. This difficulty may also apply when comparing divergence between *H. melpomene* and *H. timareta* and between *H. melpomene* and *H. cydno*.

### Sex-linked incompatibilities increase absolute Z/A divergence ratio

Despite the difficulties in directly comparing divergence on sex chromosomes and autosomes, it may be possible to detect enhanced barriers to migration on sex chromosomes (i.e. reduced *effective* migration) by comparing population pairs with different levels of *absolute* migration due to physical isolation, but that otherwise share the same common history. This can be achieved by comparing pairs of populations from the same two species that differ in their extent of geographic isolation. Indeed, previous analyses of sympatric and allopatric populations of *H. melpomene, H. cydno* and *H. timareta*, based on shared derived alleles (i.e. the ABBA-BABA test), found evidence of extensive gene flow between the species in sympatry, but with a strong reduction on the Z chromosome (Martin *et al*. 2013). Here we instead use our broad sampling scheme to investigate how patterns of sequence divergence differ with differing levels of geographic separation, and ask whether this signal can detect reduced effective migration on the Z chromosome.

Simulations show that if distance is considered a proxy for migration, reduced rates of admixture on the Z chromosome may become apparent as increased absolute Z/A divergence (*d_XY_
*(Z)/*d_XY_
*(A)) ratios over short distances, with the ratio decreasing between pairs that are geographically more isolated (Figure 7). As the effective rate of migration is reduced on the Z chromosome relative to autosomes, the *d_XY_
*(Z)/*d_XY_
*(A) ratio increases, and this increase is most pronounced when overall migration rates are high. This relation can be explained by the absolute difference in effective migration on the Z chromosome compared to the autosomes becoming smaller as overall migration decreases. While overall *d_XY_
*(Z)/*d_XY_
*(A) ratios may be influenced by ancestral population size changes, the trend should be independent from population size changes. Our widespread sampling of both clades therefore allowed us to test for reduced effective migration on the Z chromosome.

Among *H. erato* and *H. melpomene* clade populations, absolute divergence generally increases with increased distance between population pairs (Figure 7). This trend is strongest for population comparisons that are less obstructed by geographic barriers, such as among *H. erato* (mantel test: R^2^ = 0.18; p = 0.012) and *H. melpomene* (excluding Colombia; mantel test: R^2^ = 0.95; p = 0.001) populations from east of the Andes. As expected, the correlation between distance and absolute divergence is reduced by geographical barriers, such as when comparing *H. erato* (mantel test: R^2^ = 0.15; p = 0.019) and *H. melpomene* (mantel test: R^2^ = 0.55; p = 0.001) populations from Panama, east of the Andes and west of the Andes. We also observed a significant trend of increased absolute divergence with distance between populations of *H. erato* and *H. e. chestertonii* (mantel test: R^2^ = 0.44; p = 0.001), *H. melpomene* and *H. cydno* (mantel test: R^2^ = 0.69; p = 0.001) and *H. melpomene* and *H. timareta* (mantel test: R^2^ = 0.56; p = 0.001). This is consistent with gene flow among these species pairs where they are in contact. In contrast, no significant trend between absolute divergence and distance was observed between *H. erato* and *H. himera* and *H. cydno* and *H. timareta*, suggesting that these species pairs may be more strongly isolated.

We next examined the *d_XY_
*(Z)/*d_XY_
*(A) ratios and its relationship to geographic distance. If rates of admixture between populations are similar on the Z chromosome compared to the autosomes, we would not expect any relation between distance and *d_XY_
*(Z)/*d_XY_
*(A) ratios. In contrast, we observed increased *d_XY_*(Z)/*d_XY_*(A) ratios among geographically more closely located population pairs for *H. melpomene - H. timareta* (mantel test: R^2^ = 0.25; p = 0.004) and *H. melpomene* – *H. cydno* (mantel test: R^2^ = 0.35; p = 0.001) comparisons (Figure 7). Similarly, a tendency for increased *d_XY_
*(Z)/*d_XY_
*(A) ratios between *H. erato* and *H. e. chestertonii* was observed among the geographically closest comparisons, although, this was not significant (mantel test: R^2^ = 0.26; p = 0.06). Finding this trend, however, can be obscured by geographic barriers that would reduce the relation between distance and admixture. For instance, *H. e. chestertonii* comes into close contact with *H. e. venus* west of the Andes, but is geographically isolated from relatively closely located *H. erato* populations east of the Andes. Similarly, PCA analysis of the *H. melpomene* populations indicate splits between populations east of the Andes, which may reflect additional geographic barriers that do not correlate linearly with distance (Figure 1).

We also carried out a similar comparison using absolute divergence on the autosomes, which might reflect a more direct relationship with migration. Using the absolute divergence on the autosomes as a proxy for gene flow, we also find a pattern of increased Z/A divergence ratios for species pairs with known post-zygotic reproductive barriers (Figure 8). Z/A divergence ratios are significantly higher between population pairs with lower divergence values on the autosomes in *H. melpomene - H. timareta* (mantel test: R^2^ = 0.70; p = 0.001), *H. melpomene – H. cydno* (mantel test: R^2^ = 0.64; p = 0.001) and *H. erato* – *H. e. chestertonii* (mantel test: R^2^ = 0.51; p = 0.007) comparisons, but not in *H. timareta* – *H. cydno* and *H. erato* – *H. himera* comparisons. This is consistent with crosses showing that the former and not the latter pairs experience hybrid-sterility and Haldane’s Rule (McMillan *et al*. 1997; Naisbit *et al*. 2002; Salazar *et al*. 2005; Muñoz *et al*. 2010; Merrill *et al*. 2012; Sánchez *et al*. 2015) and also agrees with the previous observation of reduced shared variation between *H. melpomene* and both *H. timareta* and *H. cydno* on the Z chromosome (Martin et al. 2013). Note that our simulations suggest that to explain the trend observed in our data, there must be a very strong reduction of migration on the Z chromosome relative to the autosomes (~60% or greater).

**Figure 8.**
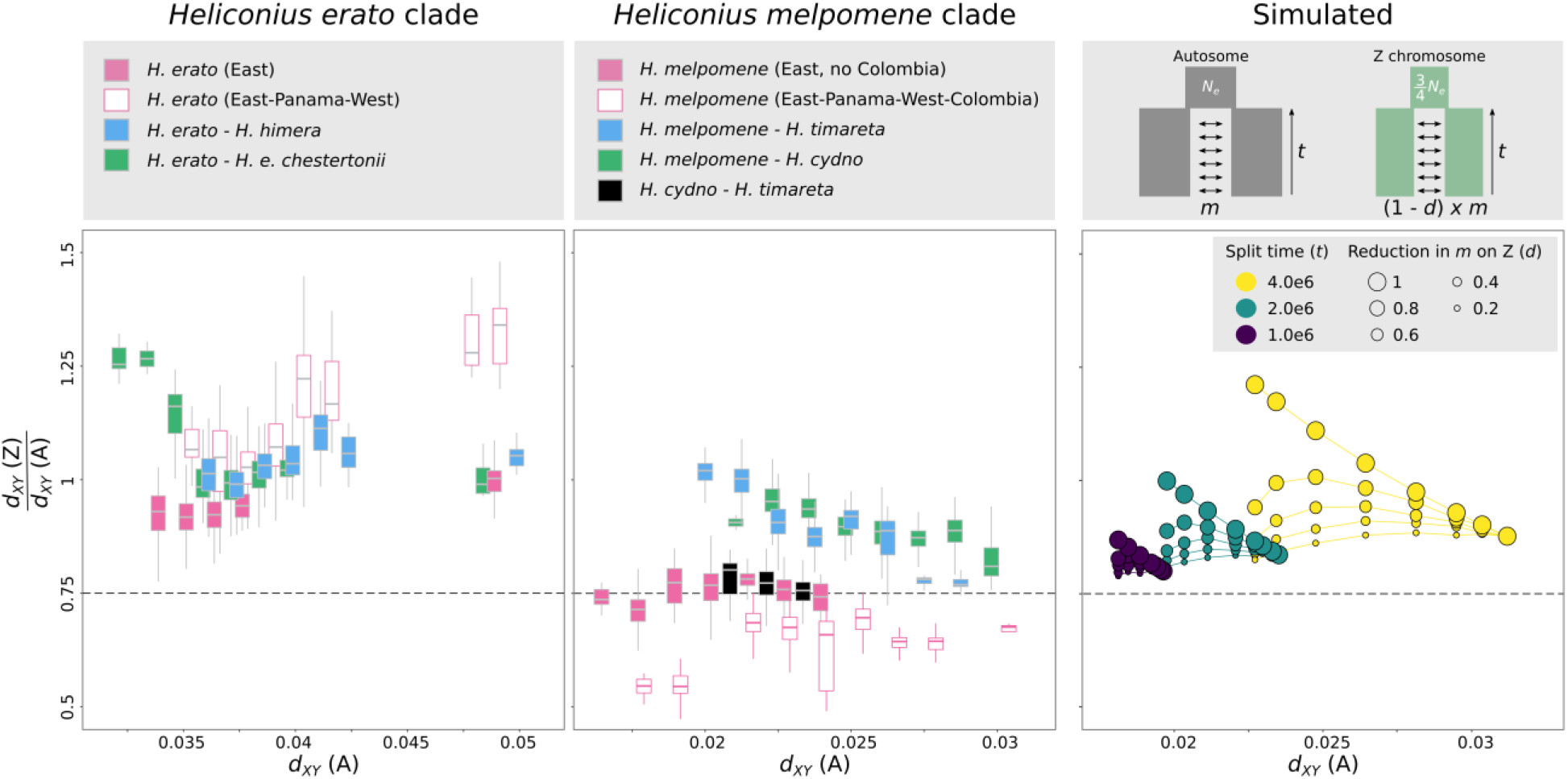
Relation between pairwise autosomal *d_XY_
* (A) and the ratio of *d_XY_
* between the Z chromosome and autosomes (*d_XY_
*(Z)/*d_XY_
*(A)) for populations of the *H. erato* and *H. melpomene* clade and simulated data. Boxplots represent pairwise measures over 1.25e-3 *d_XY_
* bins. Gray squares in the upper right panel represent the simulated populations with different rates of migration (m) expressed as a proportion of the effective population size. For the simulated data (right panels), colors indicate split times of the two populations in generations (*t*) with *N_e_
* equal to 2e6 and 1.5e6 for the autosomes and Z chromosome, respectively. Size of circles indicates proportional reduction in migration on the Z chromosome compared to the autosomes.

### Alternative factors affecting Z/A diversity ratios in Heliconius

Factors other than population size change could result in deviations from the expected Z/A diversity ratio (BOX1). In *Heliconius*, there is no empirical data on sex-biased mutation rates. While higher mutation rates on the Z could explain increased Z/A diversity ratios and increased rates of divergence (Kirkpatrick & Hall 2004; Vicoso & Charlesworth 2006; Sayres & Makova 2011), it is unlikely that closely related populations would differ in their mutation rate and that this could explain the observed variation in Z/A diversity ratios among the *Heliconius* populations. Alternatively, in *Heliconius* male-biased sex ratios have been reported in the field, which could result in increased Z/A diversity ratios. However, it has been argued that these male-biased sex ratios are most likely explained by differences in behavior resulting in male-biased captures rather than effective sex ratio differences (Jiggins 2017). Nonetheless, a *Heliconius* characteristic that could potentially amplify sex-ratio biases is that *H. erato* and *H. melpomene* clade populations are characterized by contrasting pupal-mating and adult-mating strategies, respectively (Gilbert 1976; Beltrán *et al*. 2007). Pupal-maters are largely monandrous (females mate only once), whereas adult-maters are polyandrous (Walters *et al*. 2012). Such differences in mating system could potentially result in increased variance of male reproductive success and decreased Z/A diversity ratios for monandrous mating systems (Charlesworth 2001). However, the frequency of remating in polyandrous *Heliconius* species is estimated to be only 25-30 % higher than in monandrous species (Walters *et al*. 2012) and we did not find any clear difference in Z/A diversity ratios between the pupal-mating *H. erato* and adult-mating *H. melpomene* clade populations (Figure 6). Moreover, the pattern of increased Z/A divergence that results from reduced admixture on the Z in population comparisons of geographically closely located *Heliconius* species should not be affected by sex ratio or mutation biases. Overall, in *Heliconius*, the observed variation in Z/A diversity ratios are thus likely mostly shaped by demographic changes.

### Consequences for other study systems

Similar to *Heliconius*, extensive genomic sampling is available for a number of other natural systems that have recently diverged, particularly for birds that also have ZW sex chromosomes, such as flycatchers (Ellegren *et al*. 2012), crows (Poelstra *et al*. 2014) and Darwin’s finches (Lamichhaney *et al*. 2015). In these systems, increased coalescence rates (~lineage sorting) on the Z and/or W chromosome have been accredited to the smaller effective population sizes of the sex chromosome. However, it remains unclear whether elevated measures of divergence could indicate elevated rates of between species divergence on the sex chromosomes, resulting from increased mutation or reduced admixture.

In the adaptive radiation of Darwin’s finches, there is no evidence for Haldane’s rule, nor for reduced viability of hybrids due to post-mating incompatibilities (Grant & Grant 1992) and the maintenance of isolating barriers might best be explained as resulting from ecological selection and assortative mating (Grant & Grant 2008). In crows, the divergence between hooded and carrion crows seems to be solely associated with color-mediated assortative mating even in the apparent absence of ecological selection (Randler 2007; Poelstra *et al*. 2014). Moreover, populations of both Darwin’s finches and crows can be characterized by distinct demographic histories (Lamichhaney *et al*. 2015; Vijay *et al*. 2016). Therefore, in these species, deviations in divergence measures from neutral expectation on the Z chromosome are potentially also explained by demography. In the divergence of pied and collared flycatchers, species recognition and species-specific male plumage traits are Z-linked (Saether *et al*. 2007) and female hybrids are completely sterile compared to only low levels of reduced fertility in males (Veen *et al*. 2001). In agreement with the large X-effect and disjunct rates of admixture between the sex chromosomes and autosomes, genome scans have found signals of increased relative divergence on the Z and W chromosomes (Ellegren *et al*. 2012; Smeds *et al*. 2015). The demographic history of these populations is characterized by a severe decrease in population size since their divergence (Nadachowska-Brzyska *et al*. 2013), which would influence the relative measures of divergence that have been used. Especially for the W chromosome, the reported excessive decrease of diversity and the high values of relative divergence can thus likely be partly explained by demography (Smeds *et al*. 2015). However, the excess of rare alleles (~negative Tajima’s D) on the W chromosome does contrast with these inferred demographic histories and provides support that the reduced diversity and increased *F_ST_
* measures result from selection (Smeds *et al*. 2015).

## Conclusion

The disproportionate role of sex chromosomes during speciation has been well documented based on genetic analysis, however, it is less clear how this influences patterns of divergence in natural populations. In *Heliconius*, we find much of the observed increased absolute divergence on the Z chromosome relative to neutral expectation can be explained by population size changes. This cautions against highlighting increased sex chromosome divergence as evidence for a disproportionate role in species incompatibilities or as evidence for faster X evolution. It is clear that although relative measures of divergence are most prone to demographic changes, absolute divergence measures can also be strongly influenced by population size changes. Contrary to what has been claimed, absolute measures do not therefore provide a solution to the problems inherent in using relative measures to compare patterns of divergence across genomes (Cruickshank & Hahn 2014). Despite these difficulties, we do find patterns consistent with decreased effective migration on the Z for species pairs with known post-zygotic reproductive barriers, in agreement with hybrid sterility and inviability being linked to the Z chromosome in these cases (Jiggins, Linares, *et al*. 2001; Naisbit *et al*. 2002; Salazar *et al*. 2005; Sánchez *et al*. 2015). Successfully disentangling the influence of a large-X effect and faster-X evolution on relative rates of divergence will require modeling of the demographic history of each population, including changes that may have occurred before the split of the populations. Such modelling would allow us to better contrast (i) expected within population Z/A diversity ratios with hypotheses of increased mutation rates, selective sweeps, background selection and mating system and (ii) expected between population Z/A divergence ratios with hypotheses of increased mutation rates or adaptive divergence on the Z chromosome. Additionally, our strategy of contrasting *d_XY_
*(Z)/*d_XY_
*(A) ratios with geographic distance provides opportunities for testing reduced admixture between sex chromosomes in systems for which tree-based approaches and/or crossing experiments are unfeasible.

## Materials & Methods

### sampling

We used whole genome resequenced data of a total of 109 butterflies belonging to the *Heliconius erato* clade and 115 from the *Heliconius melpomene* clade (Figure 1; Tables S1 and S2). The *H. erato* clade samples comprised fifteen color pattern forms from twenty localities: *H. e. petiverana* (Mexico, n = 5), *H. e. demophoon* (Panama, n = 10), *H. e. hydara* (Panama, n = 6 and French Guiana, n = 5), *H. e. erato* (French Guiana, n = 6), *H. e. amalfreda* (Suriname, n = 5), *H. e. notabilis* (Ecuador *H. e. lativitta* contact zone, n = 5 and Ecuador *H. e. etylus* contact zone, n =5), *H. e. etylus* (Ecuador, n = 5), *H. e. lativitta* (Ecuador, n = 5), *H. e. emma* (Peru *H. himera* contact zone, n =4 and Peru *H. e. favorinus* contact zone, n =7), *H. e. favorinus* (Peru *H. himera* contact zone, n = 4 and Peru *H. e. emma* contact zone, n = 8), *H. e. phyllis* (Bolivia, n = 4), *H. e. venus* (Colombia, n = 5), *H. e. cyrbia* (Ecuador, n = 4), *H. e. chestertonii* (Colombia, n = 7) and *H. himera* (Ecuador *H. e. emma* contact zone, n = 5 and Peru *H. e. cyrbia* contact zone, n = 4).

The *H. melpomene* clade samples comprised fourteen color pattern forms from sixteen localities (Figure 1; Tables S2). Ten populations were sampled from the *H. melpomene* clade: *H. m. melpomene* (Panama, n = 3), *H. m. melpomene* (French Guiana, n = 10), *H. m. melpomene* (Colombia, n = 5), *H. m. rosina* (Panama, n = 10), *H. m. malleti* (Colombia, n = 10), *H. m. vulcanus* (Colombia, n = 10), *H. m. plesseni* (Ecuador, n = 3), *H. m. aglaope* (Peru, n = 4), *H. m. amaryllis* (Peru, n= 10), and *H. m. nanna* (Brazil, n = 4). Three populations were sampled from the *H. timareta* clade: *H. heurippa* (Colombia, n = 3), *H. t. thelxinoe* (Peru, n = 10) and *H. t. florencia* (Colombia, n = 10). Three populations were sampled from the *H. cydno* clade: *H. c. chioneus* (Panama, n = 10), *H. c. cordula* (Venezuela, n = 3) and *H. c. zelinde* (Colombia, n = 10).

### sequencing and genotyping

Whole-genome paired-end Illumina resequencing data from *H. erato* and *H. melpomene* clade samples were aligned to the *H. erato* v1 (Van Belleghem *et al*. 2017) and *H. melpomene* v2 (Davey *et al*. 2016) reference genomes, respectively, using BWA v0.7 (Li 2013). PCR duplicated reads were removed using Picard v1.138 (http://picard.sourceforge.net) and sorted using SAMtools (Li *et al*. 2009). Genotypes were called using the Genome Analysis Tool Kit (GATK) Haplotypecaller (Van der Auwera *et al*. 2013). Individual genomic VCF records (gVCF) were jointly genotyped using GATK’s genotypeGVCFs. Genotype calls were only considered in downstream analysis if they had a minimum depth (DP) ≥ 10, maximum depth (DP) ≤ 100 (to avoid false SNPs due to mapping in repetitive regions), and for variant calls, a minimum genotype quality (GQ) ≥ 30.

### Population structure

To discern population structure among the sampled *H. erato* and *H. melpomene* clade individuals, we performed principal components analysis (PCA) using Eigenstrat SmartPCA (Price *et al*. 2006). For this analysis, we only considered autosomal biallelic sites that had coverage in all individuals.

### Population genomic diversity and divergence statistics

We first estimated diversity within populations as well as divergence between parapatric and sympatric populations in non-overlapping 50 kb windows along the autosomes and Z chromosome using python scripts and egglib (De Mita & Siol 2012). We only considered windows for which at least 10% of the positions were genotyped for at least 75% of the individuals within each population. For females, haploidi was enforced when calculating divergence and diversity statistics. Sex of individuals was inferred from heterozygosity on the Z.

*F_ST_
* was estimated as in Hudson et al. (1992), as 

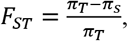

with nucleotide diversity in a population (*π_i_
*) calculated as 

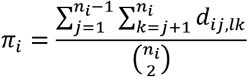

and average within population nucleotide diversity (*π_S_
*) calculated as the weighted (w) average of the nucleotide diversity (*π_i_
*) within each population *l* and *k*, as 

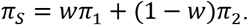

Total nucleotide diversity (*π_T_
*) was calculated as the average number of nucleotide differences per site between two DNA sequences in all possible pairs in the sampled population (Hudson *et al*. 1992), as 

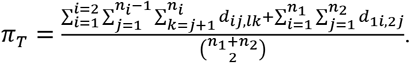

Between-population sequence divergence *d_XY_
* was estimated as the average pairwise difference between sequences sampled from two different populations (Nei & Li 1979), as 

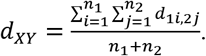

*d_a_
* was calculated as a relative measure of divergence by subtracting *d_XY_
* with an estimate of the nucleotide diversity (*π_S_
*) in the ancestral populations (Nei & Li 1979), 

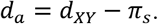

Tajima’s *D* was calculated as a measure of deviation from a population evolving neutrally with a constant size (Tajima 1989).

To overcome the problem of nonindependence between loci, estimates of the variance in nucleotide diversity (π) and Tajima’s *D* within populations along the genomes were obtained using block-jackknife deletion over 1Mb intervals along the genome (chosen to be much longer than linkage disequilibrium in *Heliconius* (Martin et al. 2013)) (Künsch 1989).

To calculate pairwise *d_XY_
* values between each individual, we subsampled the genomes by only considering genomic sites that were at least 500 bp apart and had coverage for at least one individual in each population. For the *H. erato* clade dataset, this resulted in a high coverage dataset with 322,082 and 15,382 sites on the autosomes and Z chromosome, respectively. For the *H. melpomene* clade dataset, this resulted in 335,636 and 18,623 sites on the autosomes and Z chromosome, respectively. Pairwise *d_XY_
* values between each individual were used to evaluate the relationship between absolute genetic divergence (*d_XY_
*) and geographic distance using Mantel tests (Mantel 1967). Mantel tests are commonly used to test for correlations between pairwise distance matrices and were performed using the R package *vegan* (Oksanen et al. 2016). Pairwise distances between populations were calculated from the average of the sample coordinates obtained for each population (Table S3, S4).

### simulations

To compare patterns in our data to expectations, we simulated genealogies in 50 kb sequence windows under certain evolutionary scenarios. The simulations were performed with a population recombination rate (4*N_e_r*) of 0.01 using the coalescent simulator *msms* (Ewing & Hermisson 2010). Subsequently, from the simulated genealogies, we simulated 50 kb sequences with a mutation rate of 2e-9 a Hasegawa-Kishino-Yano substitution model using *seq-gen* (Rambaut & Grass 1997).

In a first set of simulations, we considered one population that underwent a single population size change of a magnitude (*x*) ranging from 0.01 to 100 and at a certain moment backward in time (*t*). In a second set of simulations, we considered pairs of populations that were connected through migration (*m*) ranging from 0 to 1e-6 and for which migration was reduced with a factor *d* on the Z chromosome. To compare changes in the variation on autosomes and the Z chromosome, we simulated the Z chromosome as a separate population for which the effective population size was set to three-quarters that of the autosomal population.

To compare populations with a different effective population size (*N_e_
*), such as the autosomes and the Z chromosome, we expressed time in generations and migration rates as a proportion of the effective population size. Comparable to the *Heliconius* sampling, we sampled 5 individuals from each population and ran 300 replicates for each parameter combination. Pseudocode to run the *msms* command lines are provided in Tables S5. Tajima’s *D*, nucleotide diversity (*π*) and *d_XY_
* were calculated from the simulated sequences using python scripts and egglib (De Mita & Siol 2012).

## Data accessibility

Genome assemblies are available on lepbase.org. Sequencing reads are deposited in the Sequence Read Archive (SRA). See Table S1 and Table S2 for accession numbers.

## Acknowledgements

We thank the Ecuadorian Ministerio del Ambiente (No. 005-13 IC-FAU-DNB/MA), Peruvian Ministerio de Agricultura and Instituto Nacional de Recursos Naturales (201-2013-MINAGRI-DGFFS/DGEFFSS) and Autoridad Nacional De Licencias Ambientales-ANLA in Colombia (Permiso Marco 0530) for permission to collect butterflies. This work was funded by ERC grant SpeciationGenetics (339873) to CJ and NSF grant (DEB 1257689) to BC and WOM. SHM was funded by a research fellowship from St John’s College, Cambridge. CS was funded by COLCIENCIAS (Grant FP44842-5-2017). For computational resources, we thank the University of Puerto Rico, the Puerto Rico INBRE grant P20 GM103475 from the National Institute for General Medical Sciences (NIGMS), a component of the National Institutes of Health (NIH); and awards 1010094 and 1002410 from the Experimental Program to Stimulate Competitive Research (EPSCoR) program of the National Science Foundation (NSF). This work was also performed using the Darwin Supercomputer of the University of Cambridge High Performance Computing Service (http://www.hpc.cam.ac.uk/), provided by Dell Inc. using Strategic Research Infrastructure Funding from the Higher Education Funding Council for England and funding from the Science and Technology Facilities Council.

## References

Arias CF , Muñoz AG , Jiggins CD , et al. (2008) A hybrid zone provides evidence for incipient ecological speciation in *Heliconius* butterflies. Molecular ecology, 17, 4699–712.

Arias CF , Salazar C , Rosales C , et al. (2014) Phylogeography of *Heliconius cydno* and its closest relatives: Disentangling their origin and diversification. Molecular Ecology, 23, 4137–4152.

Beltran M , Jiggins C , Brower A , Bermingham E , Mallet J (2007) Do pollen feeding and pupal-mating have a single origin in *Heliconius* butterflies? Inferences from multilocus sequence data. Biological Journal of the Linnean Society, 92, 221–239.

Charlesworth B (1998) Measures of divergence between populations and the effect of forces that reduce variability. Molecular Biology and Evolution, 15, 538–543.

Charlesworth B (2001) The effect of life-history and mode of inheritance on neutral genetic variability. Genetic Research, 77, 153–166.

Charlesworth B (2012) The role of background selection in shaping patterns of molecular evolution and variation: evidence from variability on the *Drosophila X* chromosome. Genetics, 191, 233–246.

Charlesworth B , Coyne JA , Barton NH (1987) The relative rates of evolution of sex chromosomes and autosomes. The American naturalist, 130, 113–146.

Counterman B a , Ortĺz-Barrientos D , Noor M a F (2004) Using comparative genomic data to test for fast-X evolution. Evolution; international journal of organic evolution, 58, 656–660.

Coyne J , Orr H (1989) Two rules of speciation. In: Speciation and its consequences. (eds Otte D , Endler J ), pp. 180–207. Sinauer Associates, Inc., Sunderland, MA, USA.

Coyne JA , Orr HA (2004) Speciation. Sinauer Associates, Sunderland, MA, USA.

Cruickshank TE , Hahn MW (2014) Reanalysis suggests that genomic islands of speciation are due to reduced diversity, not reduced gene flow. Molecular Ecology, 23, 3133–3157.

Dasmahapatra KK , Walters JR , Briscoe AD , et al. (2012) Butterfly genome reveals promiscuous exchange of mimicry adaptations among species. Nature, 487, 94–98.

Davey JW , Barker SL , Rastas PM , et al. (2017) No evidence for maintenance of a sympatric *Heliconius* species barrier by chromosomal inversions. Evolution Letters, 1, 138–154.

Davey JW , Chouteau M , Barker SL , et al. (2016) Major improvements to the *Heliconius melpomene* genome assembly used to confirm 10 chromosome fusion events in 6 million years of butterfly evolution. G3, 6, 695–708.

De Mita S , Siol M (2012) EggLib: processing, analysis and simulation tools for population genetics and genomics. BMC Genetics, 13, 27.

Dobzhansky T (1935) Studies on hybrid sterility. II. Localization of factors in *Drosophila pseudoobscura* hybrids. Genetics, 21, 113–135.

Ellegren H , Smeds L , Burri R , et al. (2012) The genomic landscape of species divergence in *Ficedula* flycatchers. Nature, 491, 756–760.

Ewing G , Hermisson J (2010) MSMS: a coalescent simulation program including recombination, demographic structure and selection at a single locus. Bioinformatics (Oxford, England), 26, 2064–2065.

Frank SA (1991) Divergence of meiotic drive-suppression systems as an explanation for sex-biased hybrid sterility and inviability. Evolution, 45, 262–267.

Gilbert LE (1976) Postmating female odor in *Heliconius* butterflies: A male-contributed antiaphrodisiac? Science, 193, 420–422.

Gillespie JH , Langley CH (1979) Are evolutionary rates really variable? Journal of Molecular Evolution, 13, 27–34.

Grant PR , Grant BR (1992) Hybridization of bird species. Science, 256, 193–197.

Grant PR , Grant BR (2008) Pedigrees, assortative mating and speciation in Darwin’s finches. Proceedings of the Royal Society B: Biological Sciences, 275, 661–668.

Haldane JBS (1922) Sex ratio and unisexual sterility in hybrid animals. Journal of Genetics, 12, 101–109.

Hudson RR , Slatkin M , Maddison WP (1992) Estimation of levels of gene flow from DNA sequence data. Genetics, 132, 583–589.

Jablonka E , Lamb MJ (1991) Sex chromosomes and speciation. Proceedings of the Royal Society B, 243, 203–208.

Jiggins CD (2017) The ecology and evolution of Heliconius butterflies. Oxford University Press.

Jiggins CD , Linares M , Naisbit RE , et al. (2001) Sex-linked hybrid sterility in a butterfly. Evolution, 55, 1631–1638.

Jiggins CD , Mcmillan O , Neukirchen W , Mallet J , Nw L (1996) What can hybrid zones tell us about speciation? The case of *Heliconius erato* and *H. himera* (Lepidoptera: Nymphalidae). Biological Journal of the Linnean Society, 59, 221–242.

Jiggins CD , Naisbit RE , Coe RL , Mallet J (2001) Reproductive isolation caused by colour pattern mimicry. Nature, 411, 302–305.

Johnson NA , Lachance J (2012) The genetics of sex chromosomes: evolution and implications for hybrid incompatibility. Annals of the New York Academy of Sciences, 1256, E1–22.

Kirkpatrick M , Hall DW (2004) Male-biased mutation, sex Linkage, and the rate of adaptive evolution. Evolution, 58, 437.

Kronforst MR , Hansen MEB , Crawford NG , et al. (2013) Hybridization reveals the evolving genomic architecture of speciation. Cell reports, 5, 666–677.

Künsch HR (1989) The jackknife and the bootstrap for general stationary observations. The Annals of Statistics, 17, 1217–1241.

Lamichhaney S , Berglund J , Almén MS , et al. (2015) Evolution of Darwin’s finches and their beaks revealed by genome sequencing. Nature, 518, 371–375.

Lavretsky P , Dacosta JM , Hernandez-Banos BE , et al. (2015) Speciation genomics and a role for the Z chromosome in the early stages of divergence between Mexican ducks and mallards. Molecular Ecology, 24, 5364–5378.

Li H (2013) Aligning sequence reads, clone sequences and assembly contigs with BWA-MEM. arXiv, 1303.3997v1.

Li H , Handsaker B , Wysoker A , et al. (2009) The Sequence Alignment/Map format and SAMtools. Bioinformatics, 25, 2078–2079.

Mallet J , Barton NH (1989) Strong natural selection in a warning-color hybrid zone. Evolution, 43, 421–431.

Mallet J , Jiggins CD , McMillan OW (1998) Mimicry and warning colour at the boundary between races and species. In: Endless forms. Species and speciation (eds Howard DJ , Berlocher SH ), pp. 390–403. Oxford University Press, New York.

Mallet J , McMillan WO , Jiggins CD (1998) Mimicry and warning color at the boundary between microevolution and macroevolution. In: Endless Forms: Species and Speciation (eds Howard D , Berlocher S ), pp. 390–403. Oxford University Press.

Mantel N (1967) The detection of disease clustering and a generalized regression approach. Cancer Research, 27, 209–220.

Martin SH , Dasmahapatra KK , Nadeau NJ , et al. (2013) Genome-wide evidence for speciation with gene flow in *Heliconius* butterflies. Genome research, 23, 1817–1828.

McMillan WO , Jiggins CD , Mallet J (1997) What initiates speciation in passion-vine butterflies? Proceedings of the National Academy of Sciences of the United States of America, 94, 8628–8633.

Meisel RP , Connallon T (2013) The faster-X effect: Integrating theory and data. Trends in Genetics, 29, 537–544.

Mérot C , Salazar C , Merrill RM , Jiggins C , Joron M (2017) What shapes the continuum of reproductive isolation? Lessons from *Heliconius* butterflies. Proceedings of the Royal Society B: Biological Sciences, 284, 20170335.

Merrill RM , Chia A , Nadeau NJ (2014) Divergent warning patterns contribute to assortative mating between incipient *Heliconius* species. Ecology and Evolution, 4, 911–917.

Merrill RM , Wallbank RWR , Bull V , et al. (2012) Disruptive ecological selection on a mating cue. Proceedings. Biological sciences / The Royal Society, 279, 4907–4913.

Muller HJ (1942) Isolating mechanisms, evolution and temperature. Biological Symposia, 6, 71–125.

Muñoz AG , Salazar C , Castaño J , Jiggins CD , Linares M (2010) Multiple sources of reproductive isolation in a bimodal butterfly hybrid zone. Journal of Evolutionary Biology, 23, 1312–1320.

Nadachowska-Brzyska K , Burri R , Olason PI , et al. (2013) Demographic divergence history of pied flycatcher and collared flycatcher inferred from whole-genome re-sequencing data. PLoS Genetics, 9, e1003942.

Nadeau NJ , Martin SH , Kozak KM , et al. (2013) Genome-wide patterns of divergence and gene flow across a butterfly radiation. Molecular ecology, 22, 814–826.

Nadeau NJ , Whibley A , Jones RT , et al. (2012) Genomic islands of divergence in hybridizing *Heliconius* butterflies identified by large-scale targeted sequencing. Philosophical transactions of the Royal Society of London. Series B, Biological sciences, 367, 343–353.

Naisbit RE , Jiggins CD , Linares M , Salazar C , Mallet J (2002) Hybrid sterility, Haldane’s Rule and speciation in Heliconius cydno and H. melpomene. Genetics, 161, 1517–1526.

Naisbit RE , Jiggins CD , Mallet J (2001) Disruptive sexual selection against hybrids contributes to speciation between *Heliconius cydno* and *Heliconius melpomene* . Proceedings of the Royal Society B, 268, 1849–1854.

Nei M , Li WH (1979) Mathematical model for studying genetic variation in terms of restriction endonucleases. Proceedings of the National Academy of Sciences of the United States of America, 76, 5269–5273.

Oksanen J , Blanchet FG , Kindt R , et al. (2016) vegan: community ecology package. R package.

Orr HA (1996) Dobzhansky, Bateson, and the genetics of speciation. Genetics, 144, 1331–1335.

Patterson N , Richter DJ , Gnerre S , Lander ES , Reich D (2006) Genetic evidence for complex speciation of humans and chimpanzees. Nature, 441, 1103–1108.

Poelstra JW , Vijay N , Bossu CM , et al. (2014) The genomic landscape underlying phenotypic integrity in the face of gene flow in crows. Science, 344, 1410–1414.

Pool JE , Nielsen R (2007) Population size changes reshape genomic patterns of diversity. Evolution, 29, 997–1003.

Presgraves DC (2002) Patterns of postzygotic isolation in Lepidoptera. Evolution, 56, 1168–1183.

Presgraves DC (2008) Sex chromosomes and speciation in *Drosophila* . Trends in Genetics, 24, 336–343.

Price AL , Patterson NJ , Plenge RM , et al. (2006) Principal components analysis corrects for stratification in genome-wide association studies. Nature genetics, 38, 904–909.

Prowell D (1998) Sex linkage and speciation in Lepidoptera. In: Endless forms. Species and speciation (eds Howard D , Berlocher S ), pp. 309–319. Oxford University Press, New York.

Rambaut A , Grass NC (1997) Seq-Gen: an application for the Monte Carlo simulation of DNA sequence evolution along phylogenetic trees. Bioinformatics, 13, 235–238.

Randler C (2007) Assortative mating of Carrion *Corvus corone* and Hooded Crows *C. cornix* in the hybrid zone in eastern Germany. Ardea, 95, 143–149.

Sackton TB , Corbett-Detig RB , Nagaraju J , et al. (2014) Positive selection drives faster-Z evolution in silkmoths. Evolution, 68, 2331–2342.

Saether SA , Saetre G-P , Borge T , et al. (2007) Sex chromosome-linked species recognition and evolution of reproductive isolation in flycatchers. Science, 318, 95–97.

Salazar CA , Jiggins CD , Arias CF , et al. (2005) Hybrid incompatibility is consistent with a hybrid origin of *Heliconius heurippa* Hewitson from its close relatives, *Heliconius cydno* Doubleday and *Heliconius melpomene* Linnaeus. Journal of Evolutionary Biology, 18, 247–256.

Sanchez AP , Pardo-Diaz C , Enciso-Romero J , et al. (2015) An introgressed wing pattern acts as a mating cue. Evolution, 69, 1619–1629.

Sayres MAW , Makova KD (2011) Genome analyses substantiate male mutation bias in many species. Bioessays, 33, 938–945.

Slatkin M , Voelm L (1991) FST in a hierarchial island model. Genetics, 127, 627–629.

Smeds L , Warmuth V , Bolivar P , et al. (2015) Evolutionary analysis of the female-specific avian W chromosome. Nature Communications, 6, 7330.

Sperling FAH (1994) Sex-linked genes and species differences in Lepidoptera. The Canadian Entomologist, 126, 807–818.

Suomalainen E , Cook LM , Turner JRG (1973) Achiasmatic oogenesis in the *Heliconiine* butterflies. Hereditas, 302–304.

Tajima F (1989) Statistical method for testing the neutral mutation hypothesis by DNA polymorphism. Genetics, 123, 585–595.

Turelli M , Moyle LC (2007) Asymmetric postmating isolation: Darwin’s corollary to Haldane’s rule. Genetics, 176, 1059–1088.

Turelli M , Orr HA (1995) The dominance theory of Haldane’s rule. Genetics, 140, 389–402.

Turner JRG , Sheppard PM (1975) Absence of crossover in female butterflies *(Heliconius)* . Heredity, 34, 265–269.

Van Belleghem SM , Rastas P , Papanicolaou A , et al. (2017) Complex modular architecture around a simple toolkit of wing pattern genes. Nature Ecology & Evolution, 1, 52.

Van der Auwera G a. , Carneiro MO , Hartl C , et al. (2013) From fastQ data to high-confidence variant calls: The genome analysis toolkit best practices pipeline. Current Protocols in Bioinformatics, UNIT 11.10, 1–33.

Veen T , Borge T , Griffith SC , et al. (2001) Hybridization and adaptive mate choice in flycatchers. Nature, 411, 45–50.

Vicoso B , Charlesworth B (2006) Evolution on the X chromosome: unusual patterns and processes. Nature reviews. Genetics, 7, 645–653.

Vicoso B , Charlesworth B (2009) Recombination rates may affect the ratio of X to autosomal noncoding polymorphism in African populations of *Drosophila melanogaster* . Genetics, 181, 1699–1701.

Vijay N , Bossu CM , Poelstra JW , et al. (2016) Evolution of heterogeneous genome differentiation across multiple contact zones in a crow species complex. Nature Communications, 7, 13195.

Walters JR , Stafford C , Hardcastle TJ , Jiggins CD (2012) Evaluating female remating rates in light of spermatophore degradation in *Heliconius* butterflies: pupal-mating monandry versus adult-mating polyandry. Ecological Entomology, 37, 257–268.

Wolf JBW , Ellegren H (2017) Making sense of genomic islands of differentiation in light of speciation. Nature Reviews Genetics, 18, 87–100.

Wright S (1931) Evolution in mendelian populations. Genetics, 16, 97–159.

Wu C-I , Davis AW (1993) Evolution of postmating reproductive isolation: the composite nature of Haldane’s rule and its genetic bases. The American naturalist, 142, 187–212.

